# Biogeography and Validation of species limits in Caribbean Red Bats (Vespertilionidae: *Lasiurus*)

**DOI:** 10.1101/2022.02.11.479705

**Authors:** J. Angel Soto-Centeno, Camilo A. Calderón-Acevedo

**Author notes:** Corresponding author: Department of Earth & Environmental Sciences, 195 University Avenue, Rutgers University, Newark, NJ 07102, USA. (JAS-C) and (CCA).

## Abstract

Defining species limits using an integrative framework is crucial for biodiversity assessments and to maintain taxonomic stability. These approaches are robust and can be useful to also validate the status of species that are uncommon and underrepresented in biological collections. We examined the species limits and validated the taxonomic status of the Minor Red Bat (*Lasiurus minor*), an uncommon species originally described from four specimens. Our approach consisted of three independent lines of evidence combining genetic and phenotypic data. Phylogenetic analyses confirmed the uniqueness of *L. minor* compared to three other geographically and closely related Red Bat species. Furthermore, coalescent species delimitation supported the four Red Bat species hypothesis. Linear phenotypic analyses demonstrated that *L. minor* is distinct from other Red Bats despite a morphological continuum. Finally, we reassessed the diagnosability of characters used to describe *L. minor* using an objective shape analysis approach, which emphasized the support for this taxon. Based on our findings, while identification in the field could still pose a challenge, there is strong support to recognize *L. minor*. This study settles a longstanding taxonomic question and provides evidence to better understand Caribbean biodiversity.

## Introduction

Species delimitation relies strongly on the integration of diverse types of data and their complementarity to assess biologically relevant units (Carstens et al. 2013). Understanding species limits is essential to maintaining a stable taxonomy, improve the robustness of species assignment decisions, and develop accurate measures of biodiversity (Fujita et al. 2012). Because species form the basic unit of taxonomy, this process is also important for ecology and conservation biology, which inferences rely on the proper designation of species. Current methods for examining species limits provide an opportunity to also validate the limits of previously described taxa under a robust framework of integrative taxonomy (Padial et al. 2010). Thus, providing a chance to overcome the challenges presented by the revision of species that are uncommon, cryptic, and/or with few representative samples.

Caribbean Red Bats (genus *Lasiurus*) exemplify one of the most poorly documented groups across the Greater Antilles and the Bahamas (Silva Taboada 1979; Gannon et al. 2005; Speer et al. 2015; Soto-Centeno et al. 2017). Two taxonomic hypotheses of Caribbean Red Bats have been proposed based primarily on limited phenotypic or genetic data. Some authors have considered Caribbean Red Bats a complex of three species (sensu Miller 1931), i.e. *Lasiurus degelidus* (Jamaica), *L. minor* (the Bahamas, Hispaniola, and Puerto Rico), and *L. pfeifferi* (Cuba); yet others considered all Caribbean Red Bats as a single species attributed to *L. borealis* (sensu Koopman et al. 1957). Studies have confirmed the unique taxonomic identities of the Cuban and Jamaican Red Bats confirming the original designation of Miller (1931) (Baker et al. 1988; Baird et al. 2015).

The Minor Red Bat (*Lasiurus minor*) has received considerable attention regarding its taxonomic status. This species was described by Miller (1931) on the basis of four skulls retrieved from owl pellets collected in Voute l’Église cave in Jacmel, Haiti. The skulls of *L. minor* were characterized as “noticeably smaller” than those of its congeners from Cuba (*L. pfeifferi*) and Jamaica (*L. degelidus*) with variation in four characters (Miller 1931). These four characters, when compared with *L. borealis*, were a braincase “more rounded when viewed from above, and more flat-topped when viewed from behind; lacrimal ridge and tubercle poorly developed; upper cheekteeth essentially like those of *L. borealis borealis* except that PM^4^ is smaller” (Miller 1931, p.410). Two Red Bat specimens collected on Cat Island and New Providence, the Bahamas, were also conferred to *L. minor*, extending the distribution of this species (Allen and Sanborn 1937). In a revision of Red Bats from the Bahamas, Koopman et al. (1957) agreed with Allen and Sanborn (1937) that the Bahamian species are best conferred to *L. minor*. Notwithstanding, they concluded that given the paucity of the available material and the lack of suitable diagnostic characters, *L. minor* should be considered a “race” of *L. borealis* (Koopman et al. 1957, p.168). The first Red Bat report from Puerto Rico described a specimen with general features of the skull most closely resembling the description of *L. minor* from Haiti (Starret and Rolle 1962). Yet the most recent record reported of a reproductive Red Bat female for the island followed the taxonomic proposal of Koopman et al. (1957) of *L. borealis minor* (Rodríguez-Durán 1999). Still, despite that no recent or thorough revisions to test the species limits have been made, the taxon *Lasiurus minor* (Miller 1913) remains valid (Simmons 2005; Lim et al. 2017; Simmons and Cirranello 2020).

In this study, we provide independent evidence to validate the species limits of the Minor Red Bat (*Lasiurus minor*, Vespertilionidae). Specifically, we tested the hypothesis that *L. minor* constitutes a unique lineage at the species level that is distinct both genetically and morphologically from other Red Bats that are closely related and geographically adjacent. We used a species validation approach to provide additional support to the recognition of this taxon under an integrative and statistically robust framework.

## Materials and Methods

### Gene sequences

We compiled all available sequences for a fragment of the mitochondrial cytochrome oxidase I (COI) gene encompassing 657 bp in length. Sequences were obtained from already published records of Red Bats (*Lasiurus blossevillii, L. borealis, L. frantzi, L. minor, L. pfeifferi*, and *L. seminolus*) included in (Streicker et al. 2010; Clare et al. 2011; Baird et al. 2015; Lim et al. 2017). Identical sequences were excluded from all analyses and the resulting set of sequences (N = 50) was aligned using the Muscle algorithm in Geneious Prime v2020.1.2 (Biomatters Ltd.). A single COI sequence of the big brown bat (*Eptesicus fuscus*) from Virginia, USA was included as outgroup in the phylogenetic analysis. Alignment file used in this study is available in fasta format and archived in http://dx.doi.org/10.17632/79m4fj8rdx.1.

### Phylogenetic inference

To estimate the phylogenetic relationships and specifically determine the position of *L. minor* among all other Red Bats, we used a Maximum Likelihood approach implemented in RAxML-NG (Kozlov et al. 2019). This analysis consisted of a GTR+G model of substitution and run over 50 independent tree searches using 25 random and 25 parsimony-based starting trees to produce the best scoring topology. This analysis was repeated in three independent inference runs and loglikelihoods were compared by computing the topological Robinson-Foulds (RF) distances to ensure a thorough exploration of the likelihood surface (Robinson and Foulds 1981). Branch support was inferred by computing 1000 MRE-based bootstrap replicates (Pattengale et al. 2010) where an auto MRE cutoff of 0.03 was used to automatically assess if sufficient replicates were performed. The summarized branch support was computed as Transfer Bootstrap Expectation (TBE; Lemoine et al. 2018) metrics from all bootstrap replicates plotted onto the best topology. TBE can reliably recover support because it is based on the minimum transfer distance between a given branch and any other branch in the bootstrap replicate tree (Lemoine et al. 2018).

### Species delimitation

Our primary goal was to examine the validity and species limits of *L. minor* in relation to other Red Bats of geographically close species from the Caribbean and the Southeastern United States. Thus, following the same approach as above, and considering the conclusion of Koopman et al. (1957), we developed a concise alignment only including unique sequences (N = 33) of our Red Bat focus group: *L. borealis, L. minor, L. pfeifferi* and *L. seminolus*. Species limits within these Red Bats were examined using two independent approaches with the single mitochondrial COI locus.

First, we analyzed limits using the multi-rate Poisson Tree Process, a tree-based method that can accommodate different rates of coalescence within clades and is optimized to handle barcoding loci (mPTP; Kapli et al. 2017). The mPTP analysis was run in the server https://mcmc-mptp.h-its.org/mcmc and used a non-ultrametric tree for the focal taxa produced in RAxML-NG. The MCMC runs included 10 × 10^6^ generations that were sampled every 10 thousand after a 10% burn-in. Three separate analyses were set with different starting delimitation models: null model (i.e. all taxa considered as one species), maximum likelihood model (i.e. MLE based delimitation), and random model (i.e. random delimitation). In each analysis, the option *--multi* was used to include the intra-specific differences among rates of coalescence with a minimum branch length of 0.001.

Second, we also tested species limits using the Bayesian coalescent-based approach BPP v4.2.9 (Yang and Rannala 2014) following the guidelines of Flouri et al. (2018). We ran an initial species tree estimation (A01 analysis) using the fixed maximum likelihood tree as a guide, a diffuse prior for *θ* in which α = 3 and β was estimated from the mutation rate for the available COI sequences. The diffuse prior of (included α = 3 and adjusting β ranging from 0.02 to 0.002 on multiple test runs to ensure convergence of the mean onto the divergence estimate of *L. borealis* in Baird et al. (2015). Additionally, we used an unguided delimitation analysis (A11 analysis) to compare the consistency of the resulting species limits. Each run consisted of an MCMC chain of 2 × 10^6^ generations, sampling every second generation with a 10% burn-in. Species delimitation was tested in five independent runs to ensure the reliability of results. Convergence was determined by examining the loglikelihood values of each run using Tracer v1.7 (Rambaut et al. 2018).

### Specimen data

To examine phenotypic differences among Red Bat species, we obtained 147 crania and dentaries as loan from *L. borealis* (N = 78), *L. pfeifferi* (N = 1), and *L. seminolus* (N = 69) deposited in the Department of Mammalogy at American Museum of Natural History (Supplementary Data S1). Recent specimens of *L. minor* are notoriously underrepresented in museum collections that are broadly available to the scientific community, in part due to the rarity of this species and the paucity of systematic collection efforts. Thus, our sample for this species constituted 88 fossil elements collected from Trouing Jean Paul (TJP), a limestone sinkhole cave located in Parc National La Visite, Massif de la Selle, Haiti (18.33°N, - 72.28°W) and described in (Soto-Centeno et al. 2017). This locality is about 50 km east of the type locality for *L. minor*. TJP is a high elevation site (∼1825 m) excavated in February 1984 by a field team led by Charles A. Woods. Relevant documentation associated with the fossil excavations were obtained by C. A. Woods at the Florida Museum of Natural History, University of Florida (UF). These fossils were loaned to us for identification and study by the UF Division of Vertebrate Paleontology, where permits and field notes are archived. Radiocarbon date analysis of the fossil *L. minor* indicated that these specimens ranged from 1690–570 cal. yr. BP (2σ; Soto-Centeno et al. 2017). The total number of specimens measured from the focus Red Bat species was 236.

### Morphology: machine learning classification models

We measured seven cranial characters to the nearest 0.01 mm from 226 Red Bat specimens (see Specimen Data) using digital calipers (Mitutoyo, Japan). Each measurement was taken three times to account for error. The characters examined included post orbital width, the premolar to molar distance, the distance between proximate ridge of the nasal to the distal point of the occipital lobe, the length of the palate, the condylobasal length, the length of the narrowest point of distal surfaces of the pterygoid plates, and the distance between the anterior most point of the glenoid fossa to the origin of the masseter muscle (see Jacobs 1996). Some measurements could not be recorded because several *L. minor* fossils were fragmented. Thus, to maximize our sample size, we partitioned the data per species and used the multivariate imputation by chained equations approach in the R package *mice* (Van Buuren and Groothuis-Oudshoorn 2011) ensuring that all measurements in each species did not exceed 40% missing data before imputation (Penone et al. 2014). Final dataset used in this study is available as a text file and archived in http://dx.doi.org/10.17632/79m4fj8rdx.1.

We used these seven cranial measurements to test the hypothesis that *L. borealis, L. minor*, and *L. seminolus* form diagnosable phenotypic groups. *Lasiurus pfeifferi* was not included in this and subsequent morphological analyses because the N = 1 precluded examination of variability and violated the assumptions of the model. To examine phenotypic divergence, we built a classification model under a supervised machine learning approach of Linear Discriminant Analysis (LDA) in the R packages *caret* (Kuhn 2020) and *MASS* (Venables and Ripley 2002). The classification model was trained using a 75% random data partition and then tested using the remaining 25% of the data implementing a k-fold cross validation approach of five replicates. We then computed a confusion matrix to calculate model accuracy, or how well the classifier assigned each species to its correct group, and then evaluated whether the overall accuracy rate was greater than the no-information rate (Kuhn and Johnson 2013). Phenotypic limits were examined on a two-dimensional plot of the first two linear discriminants.

### Morphology: elliptical Fourier shape descriptors and gaussian models

We tested the validity and ability of the characters proposed by Miller (1931) to distinguish *L. minor* from *L. borealis* and *L. seminolus* using an objective approach for shape quantification. To quantify the shapes of these four characters (see Introduction), we traced the contour of each from digital photographs of the skull using Adobe Photoshop (CC 2019) and ImageJ (Schneider et al. 2012). The specific contours were: 1) the braincase from a dorsal view, following the contour of the skull from the narrowest post orbital point around the braincase, including the frontal, parietal, squamosal, and post parietal but excluding the occipital condyle, the mastoid process, and zygomatic arch (N = 24 *L. borealis*, 29 *L. minor*, and 18 *L. seminolus*); 2) the posterior of the skull from the left mastoid process, reaching the top of the skull and descending into the right mastoid process (N = 27 *L. borealis*, 39 *L. minor*, and 21 *L. seminolus*); 3) the contour of the prefrontal and frontal bone margins around the lacrimal ridge from a dorsal view (N = 26 *L. borealis*, 37 *L. minor*, and 19 *L. seminolus*); and 4) the shape of the last upper premolar (PM^4^; N = 28 *L. borealis* and 38 *L. minor*) from a lateral view (Supplementary Data SD2).

Bitmap formatted images of the contours were analyzed in the software SHAPE v1.3 (Iwata and Ukai 2002). Each was transformed into chain code, assigning a string of code that represented the contour of every individual image of the diagnostic characters of *L. minor*. We used each contour to create a harmonic or elliptical Fourier descriptor (EFDs) series. This was used to quantify the shape of the diagnostic characters of *L. minor* in comparison to *L. borealis* and *L. seminolus*. The harmonics depict different coordinates or descriptors of a shape, which were used as input for a Principal Component Analysis (PCA) to determine the position of each species in morphospace. We allowed SHAPE v1.3 to select the number of effective principal components that best explained shape variation within our sample.

PCA scores of the EFD series represent morphological characters of biological significance. Therefore, we used the scores to examine the ability of each character to discriminate between the three Red Bat species. We fit gaussian mixture models (GMMs; McLachlan and Peel 2000; McLachlan et al. 2019) on those PCA scores. GMMs provide a statistical framework that extend ordination analyses like PCA. This method uses the mixture of probability distributions underlying a continuous character dataset. It allows to examine the combination of distributions that better explain the phenotypic variation in a mixture of components (i.e. morphological clusters) present in the dataset, and can be used as a guided discriminant analysis (Fraley and Raftery 2002). The parameters of GMMs include means and variance-covariance matrices, which describe the phenotypes of groups detected among specimens depending on the normal distributions underlying the Principal Components derived from the SHAPE analysis. This flexible and objective framework allowed to test the discrimination power of the original characters described by Miller (1931) in a model based discriminant analysis using the MclustDA function in the R package *Mclust* v5.4.7 (Scrucca et al. 2016). Output files from SHAPE v1.3 are available as native files archived in http://dx.doi.org/10.17632/79m4fj8rdx.1.

## Results

### Phylogenetics and species limits

Phylogenetic analysis of the mitochondrial COI gene clarified relationships among Red Bats from the Caribbean and the Southeastern United States (Fig. 1). The best topology resulted in a final -ln = -2789.362. Each species in our focus group (i.e. *L. borealis, L. minor, L. pfeifferi*, and *L. seminolus*) formed well supported monophyletic clades. COI was not variable enough to resolve the position of *L. frantzii*, although this taxon was outside the scope of the study. For our focus group, uncorrected p genetic distances within species were below 1%. In contrast, the COI gene between geographically close species shows at least 5% divergence in *L. pfeifferi* from Cuba and *L. seminolus* from Southeast US, and up to 9% divergence between *L. pfeifferi* and the geographically nearby *L. minor* from Dominican Republic (see Supplementary Data SD3).

**Figure 1.**
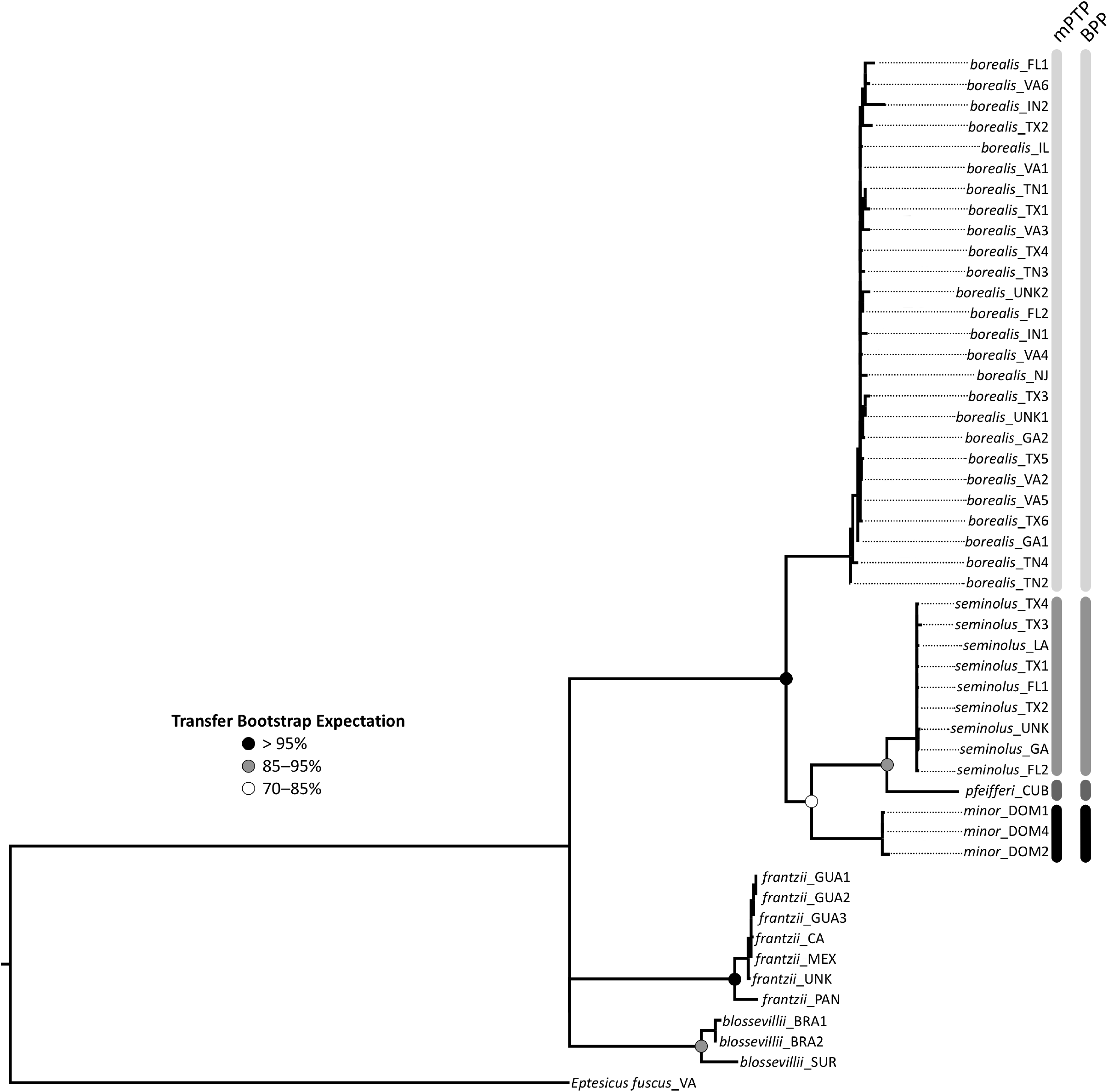
Maximum Likelihood phylogeny of mitochondrial COI sequences of Red Bats (genus *Lasiurus*). Colored circles at nodes represent Transfer Bootstrap Expectation (TBE) values estimated in RaxML-NG. Each vertical bar corresponds to summarized results of mPTP and BPP species delimitation analyses based on the concise alignment. Both mPTP and BPP results agree on four species, supporting the validity of *Lasiurus minor*.

The species tree analyses confirmed that our focus group consists of four species of Red Bats (Fig. 1). The three independent methods used as starting delimitation in the mPTP analyses (i.e. null model, maximum likelihood, or random) all strongly inferred the four species tree. Across all combinations of prior settings, BPP analyses rejected the null hypotheses that the four Red Bats in our focus group belong to a single species. The guided (A10) and unguided (A11) analyses in BPP strongly supported that our focus group consists of four species. However, these analyses could not specifically resolve the position of *L. minor* relative to its conspecifics (Table 1). Overall guided and unguided analyses in BPP resulted in three alternative topologies, including one identical to Fig. 1, with a combined best posterior probability ranging from 0.982 to 0.997 (Table 1).

**Table 1.**
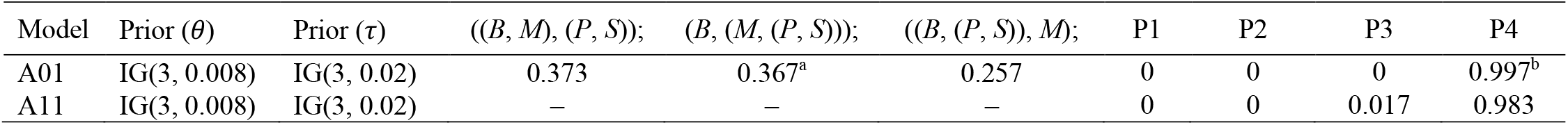
Outline of priors and posterior probability results for species delimitation analyses of Red Bats in BPP. Two main analyses were run using a fixed guide tree (A01) or unguided (A11). Two different prior scenarios were examined under the different analyses based on population size (inverse gamma *θ* = *α*, β) and divergence time differences (inverse gamma *τ* = *α*, β). In the case of population size, β was estimated from the mutation rate of COI gene sequences and then fixed for subsequent tests. In the three alternative topologies *B* = *L. borealis, M* = *L. minor, P* = *L. pfeifferi*, and *S* = *L. seminolus*. In A01 analyses, the initial topology is indicated by ^a^ and it matched the relationships obtained from best supported Maximum Likelihood tree (see Fig. 1). The combined posterior probability of the best three out of ten alternative topologies is indicated by ^b^. P1–4 indicate the posterior probabilities of species delimitation from one to four species.

### Morphology

The percent group separation achieved by the linear discriminants were 86.9 for LD1 and 13.1 for LD2. The machine learning LDA classifier of phenotypic limits in Red Bats had an overall accuracy of 77.3% (95% CI: 70.3–81.8%), which was significantly greater than the no information rate (*P* < 0.005). The greatest extent of phenotypic overlap was observed between *L. borealis* and *L. seminolus* with 22% and 28% incorrectly assigned among these species, respectively (Fig. 2). The classifier achieved better separation of *L. minor* from *L. borealis* and *L. seminolus*. Notwithstanding, 10% and 5% of *L. minor* were incorrectly assigned to *L. borealis* and *L. seminolus*, respectively (Fig. 2).

**Figure 2.**
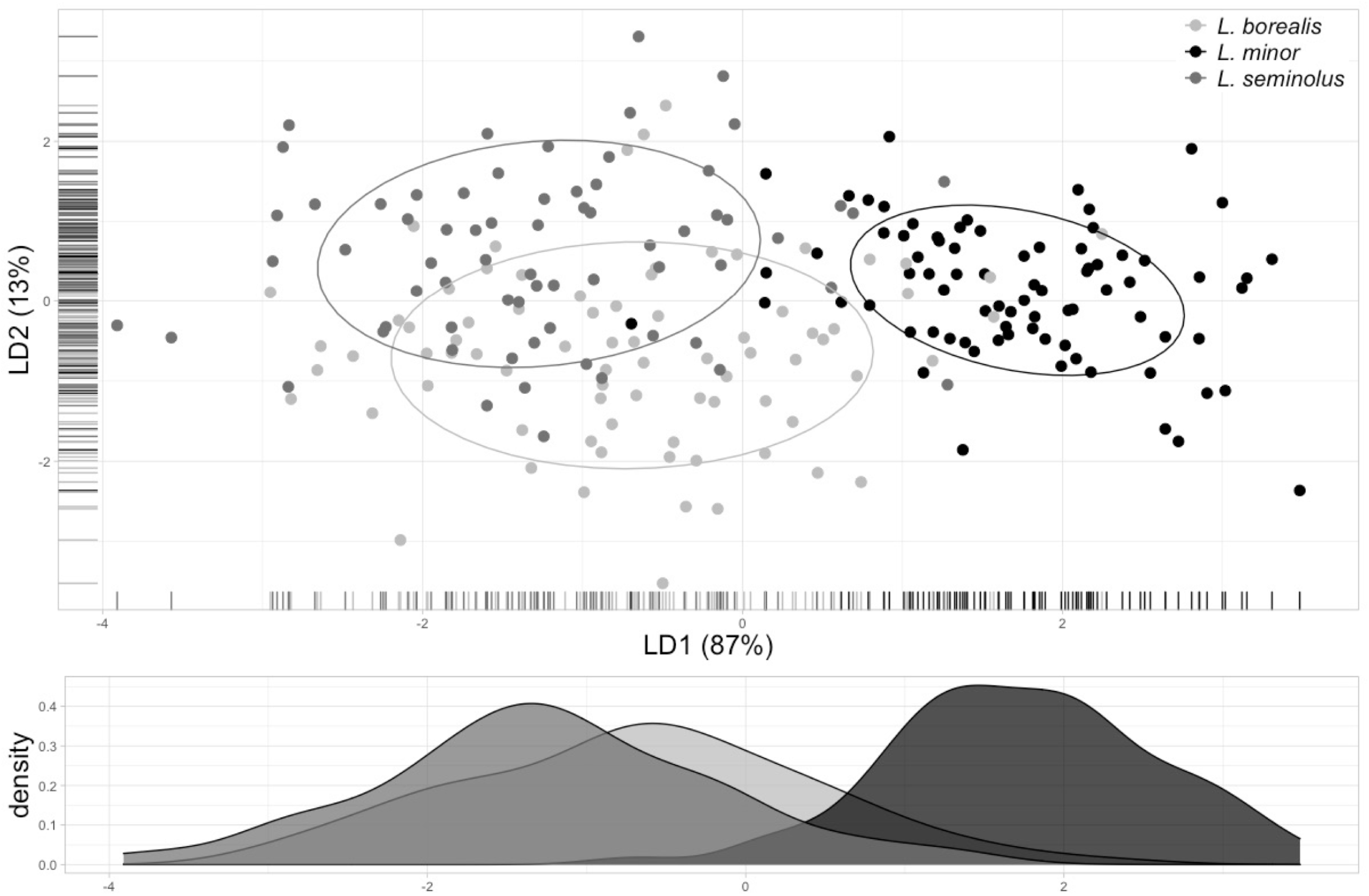
Results from the machine learning LDA classifier of phenotypic limits in Red Bats (genus *Lasiurus*). Overall accuracy of the model = 77.3% (95% CI: 70.3–81.8%). Percent group separation described for the first linear discriminant (LD1) was 86.9 and 13.1 for the second linear discriminant (LD2). Solid lines represent 68% data ellipses to visualize phenotypic overlap among species. LD1 density values plotted for aid in visualization on the x-axis where *L. minor* separates from *L. borealis* and *L. seminolus. L. pfeifferi* was not included in this analysis due to low sample size.

The PCAs of four different character shapes showed that *L. minor* occupies a section of morphospace that overlapped with some *L. borealis* or *L. seminolus* (Fig. 3A, B, D). In contrast, the shape of the lacrimal ridge showed little overlap with other species and a more cohesive cluster of *L. minor* (Fig. 3C). Despite some overlap in morphospace with *L. borealis* and *L. seminolus, L. minor* tended to occupy a section of morphospace with rounder and flatter skulls and last upper premolar with a less pronounced hypoconal basin. These, coupled with the poorly developed lacrimal ridge, supported the use of these diagnostic characters by Miller (1931). The gaussian model discriminant analysis did reliably identify and discriminate between all three Red Bat species. The roundness and flatness of the skull showed a classification error rate of 80% in GMM. The last upper premolar and the lacrimal ridge were more diagnosable characters, with an error rate of 90% and 97%, respectively.

**Figure 3.**
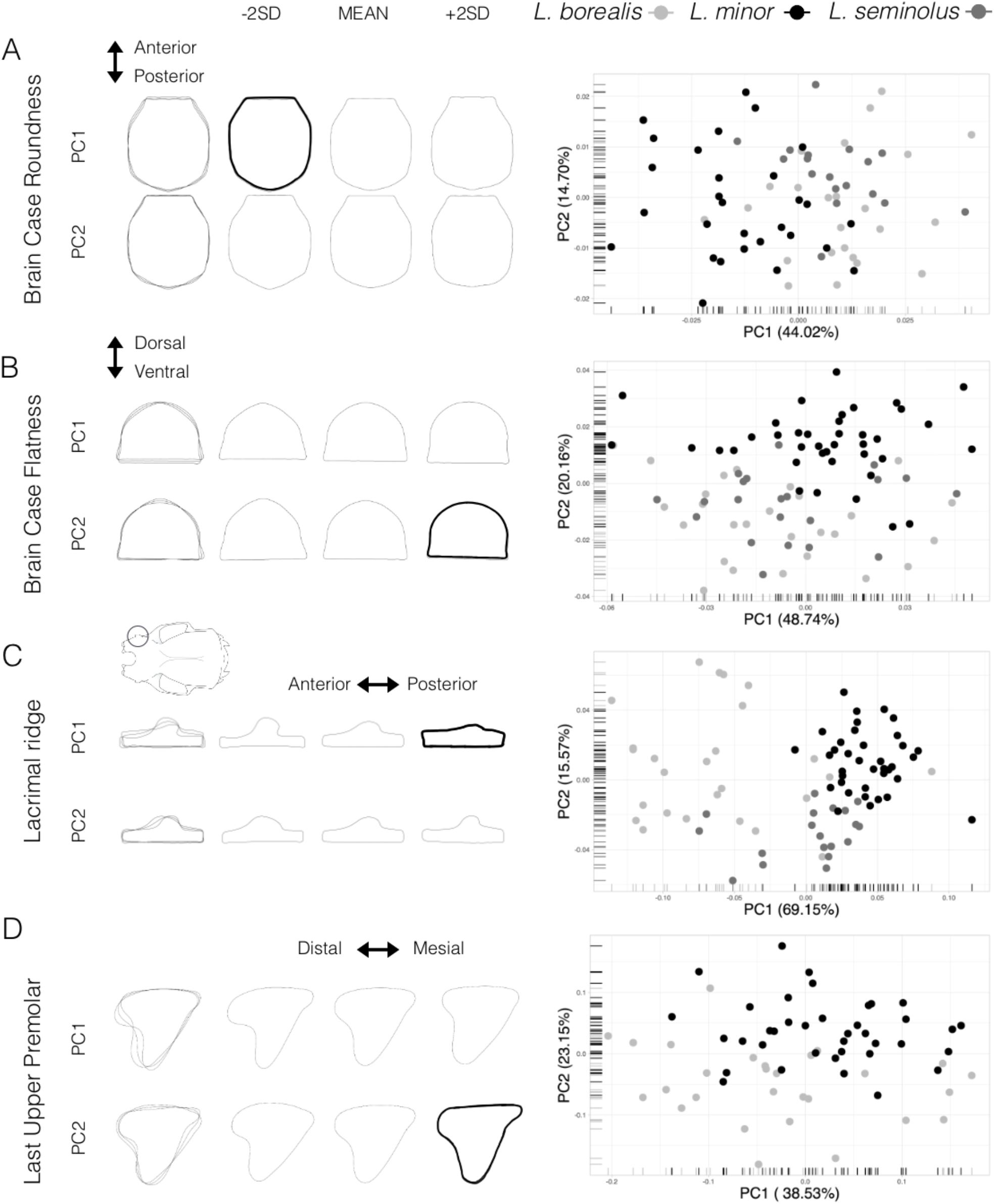
Shape variation of Red Bat diagnostic characters proposed by Miller (1931) and PCA scatterplots showing the morphospace occupied by *L. borealis, L. minor*, and *L. seminolus*. A) dorsal view of the skull representing the roundness of the braincase, B) posterior view of the skull representing the flatness of the braincase, C) dorsal views of the skull showing structural differences of the lacrimal ridge, and D) lateral view of the last upper premolar (PM4). The left panel shows the range (i.e. mean ±2 SD) for each shape examined under the first two principal components. Shape highlighted in black indicates the closest shape estimate representing *L. minor*. For details of image processing see Supplementary Data SD2.

## Discussion

A major principle of integrative taxonomy is to provide an objective framework using multiple lines of evidence to examine patterns of species divergence to evaluate taxonomic status. The use of different criteria to validate species (e.g. lineage monophyly and morphological distinctiveness) can resolve species limits by providing the evidence necessary to reject or support existing taxonomic hypotheses (Coyne and Orr 2004). Furthermore, these methods can aid in untangling the complexity of different evolutionary histories that contribute to differentiation (de Queiroz 2005). Our goal was to provide a robust framework to validate the hypothesis that the Minor Red Bat (*Lasiurus minor*) constitutes a unique lineage at the species level that is distinct from other Red Bats. Thereby, providing taxonomic stability and clear the disagreements on this Caribbean Red Bat group (Miller 1931; Allen and Sanborn 1937; Koopman et al. 1957; Starret and Rolle 1962; Rodríguez-Durán 1999; Gannon et al. 2005; Simmons 2005; Cláudio 2019; Simmons and Cirranello 2020) The results of our study based on an integrative approach of multiple data types and analyses showed support for the recognition of *L. minor* (Miller 1931).

Two recent studies examined Red Bats in a phylogenetic context. Baird et al. (2015) confirmed the phylogenetic relationships of *L. borealis* as an early divergent lineage sister to *L. pfeifferi* and *L. seminolus*. However, no samples of *L. minor* were available in that study to examine the phylogenetic placement this Caribbean Red Bat lineage. Lim et al. (2017) in a broad phylogenetic survey of the bats of Dominican Republic produced the first sequences of *L. minor*. All four samples originate from Parque Nacional Armando Bermúdez, near Pico Duarte, the highest peak in the entire Caribbean. These samples of *L. minor*, however, were only discussed in the context of interspecific comparisons of genetic variation among all bats in the Dominican Republic (Table 2 in Lim et al. 2017).

**Table 2.**
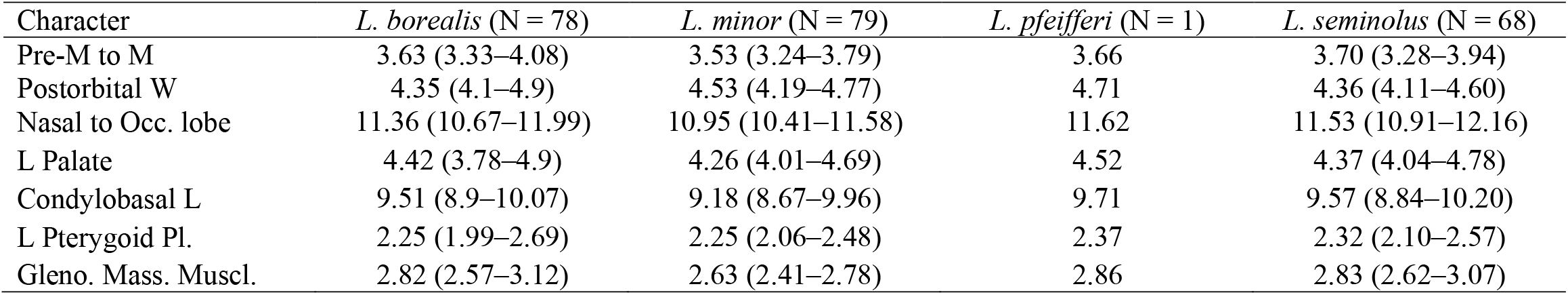
Measurements of four species of Red Bats (genus *Lasiurus*) examined in this study. Measurements represented as means with ranges in parenthesis. Sample sizes noted in parenthesis next to each species. Character abbreviations are: Pre-M to M = Pre-molar to Molar distance, Postorbital W = post-orbital width, Nasal to Occ. Lobe = proximate ridge of the nasal to the distal point of the occipital lobe, L Palate = length of palate, Condylobasal L = condylobasal length, L Pterygoid Pl. = length of narrowest point of distal surfaces of pterygoid plates, and Gleno. Mass. Muscl. = distance between the glenoid fossa to the origin of the masseter muscle.

| Character | *L. borealis* (N = 78) | *L. minor* (N = 79) | *L. pfeifferi* (N = 1) | *L. seminolus* (N = 68) |
| --- | --- | --- | --- | --- |
| Pre-M to M | 3.63 (3.33–4.08) | 3.53 (3.24–3.79) | 3.66 | 3.70 (3.28–3.94) |
| Postorbital W | 4.35 (4.1–4.9) | 4.53 (4.19–4.77) | 4.71 | 4.36 (4.11–4.60) |
| Nasal to Occ. lobe | 11.36 (10.67–11.99) | 10.95 (10.41–11.58) | 11.62 | 11.53 (10.91–12.16) |
| L Palate | 4.42 (3.78–4.9) | 4.26 (4.01–4.69) | 4.52 | 4.37 (4.04–4.78) |
| Condylbasal L | 9.51 (8.9–10.07) | 9.18 (8.67–9.96) | 9.71 | 9.57 (8.84–10.20) |
| L Pterygoid Pl. | 2.25 (1.99–2.69) | 2.25 (2.06–2.48) | 2.37 | 2.32 (2.10–2.57) |
| Gleno. Mass. Muscl. | 2.82 (2.57–3.12) | 2.63 (2.41–2.78) | 2.86 | 2.83 (2.62–3.07) |

Herein, the phylogenetic and species delimitation analyses confirm the uniqueness of *L. minor* and support the four species hypothesis (i.e. *L. borealis, L. minor, L. pfeifferi*, and *L. seminolus*; Fig. 1). Among these four focal taxa, our phylogenetic analysis showed *L. borealis* as an early divergent lineage sister to the *L. minor, L. pfeifferi*, and *L. seminolus* clade. Contrary to the suggestion of Koopman et al. (1957) to place *L. minor* as a “race” of *L. borealis*, these data indicate with high confidence that *L. minor* is a sister clade to *L. pfeifferi* and *L. seminolus* (Fig. 1). The placement of *L. minor* within this clade, however, was recovered with moderate support.

The BPP analyses resulted in three alternative topologies with almost equal posterior probability support (Table 1). This indicates that the COI locus used has enough information to validate the species but only weak phylogenetic information to infer the specific placement of *L. minor*, a result consistent with the moderate transfer bootstrap expectation value of the phylogenetic analysis. We recognize that the idiosyncratic history of a single locus may not fully account for the evolutionary history of a species (Collins and Cruickshank 2012; Alvarado-Serrano and Hickerson 2016). Nevertheless, as shown in simulation studies (Zhang et al. 2014), the error rate of inferring species delimitation by BPP is low, even when using a single locus. Our estimated guide tree (see Phylogenetic Inference in Methods) was robust and able to overcome the high false positive rates of species delimitation estimates associated when using a random guide tree (Leaché and Fujita 2010; Yang and Rannala 2014; Zhang et al. 2014). Finally, we confirmed the reliability of the delimitation results in two ways. First, we independently examined the sensitivity of our analysis using guided and unguided analyses in BPP. Second, we implemented three different starting delimitation models in mPTP, a single locus species delimitation approach that is conservative and likely to represent the true species clusters (Blair and Bryson 2017; Kapli et al. 2017). All species delimitation analyses strongly supported the four species tree hypothesis (Fig. 1; Table 1). Combined, these steps provided phylogenetic confirmation of *L. minor* as a unique lineage separate from *L. borealis* and established a platform to further examine the validity of this species from an independent phenotypic perspective.

The *L. minor* specimens available to us for examination were fossils (1690–570 cal. yr. BP) and represent the largest sample of Minor Red Bats available to date (Soto-Centeno et al. 2017). A conservative total of seven characters was used to develop a phenotypic dataset of cranial linear measurements because some fossils had missing fragments. We organized these characters into species groups a priori based on the genetic evidence discussed above. The LDA classification model, thus, was explicitly designed to test whether these independent characters could explain the species limits observed. Some authors understandably regarded the paucity of phenotypic variation as a cautionary note to make taxonomic decisions about *L. minor* (Koopman et al. 1957; Gannon et al. 2005; Cláudio 2019). The LDA classification model we present uncovered that phenotypic variation among *L. borealis, L. minor*, and *L. seminolus* represents a continuum instead of the typically assumed hierarchical morphological structure (see Zapata and Jiménez 2012). Despite that continuity, phenotypic separation of *L. minor* from other Red Bats was observed along the post-orbital width and the pre-molar to molar distance represented in LD1 and LD2 (Fig. 2). It is important to note that our LDA classifier (average accuracy = 77.3%) strongly discriminated *L. minor* with a precision rate of 0.94, whereas *L. borealis* and *L. seminolus* obtained a precision rate of 0.65 and 0.70, respectively. This emphasizes the phenotypic distinction of *L. minor* despite the lack of morphological variation noted by other authors (Koopman et al. 1957). In contrast, *L. borealis* and *L. seminolus* showed greater phenotypic similarity as confirmed by their broader overlap (Fig. 2). If additional characters with better phenotypic signal were chosen (e.g. from recent specimens), we would expect the separation of *L. minor* to become better defined.

Species descriptions, diagnosis, and delimitation are often proposed using a few specimens and characters (e.g. Miller 1931; and many others). The interpretation of diagnosable morphological characters can include substantial subjectivity, especially in species that span a broad geographic range and environmental conditions (Cadena et al. 2017). We avoided this pitfall by combining the quantification of character shapes under a modelling approach that accounts for all components that describe such shapes (McLachlan and Peel 2000; Iwata and Ukai 2002; Scrucca et al. 2016; McLachlan et al. 2019). These methods are replicable and free of bias from subjective interpretations of investigators; thus, adding a more robust and stable examination character diagnosability. Under this framework, we reevaluated the validity of the four diagnosable characters proposed by Miller (1931) to delimit *L. minor* from other closely related Red Bats. The morphological variation captured in our SHAPE analysis and used as input for our GMMs provided a robust way to test the discriminant efficacy of skull and tooth shapes. Our results showed that the probability distributions of these principal components have different degrees of effectiveness in discriminating the different morphological clusters into each Red Bat species. Two of these characters, the roundness and flatness of the braincase, had little discrimination power (Fig. 3A, B). In contrast, we showed that the shape of the lacrimal ridge and the shape of the last upper premolar (PM^4^) were more diagnosable and potentially useful for biologists to discriminate *L. minor* from *L. borealis* or *L. seminolus* (Fig. 3C, D).

Based on the results of character shapes, *L. minor* can be diagnosed by having a reduced lacrimal ridge and a PM^4^ that lacks a distinctive hypoconal basin (Fig. 3C, D). However, we caution that due to the breadth of variability of these characters observed among *L. borealis, L. minor*, and *L. seminolus*, morphological identification in the field could still be challenging. There are few places where Red Bats in the Caribbean occur in high densities. Therefore, field comparisons of these characters based on a single or a few captured individuals may lead to misidentifications, particularly in geographic areas where all species may occur in sympatry (e.g. the Bahamas). We believe that this character variability and the paucity of specimens examined from an area of sympatry led Koopman et al. (1957) to the conclusion that without appropriate evidence it is best to synonymize *L. minor* with *L. borealis*.

Integrative analyses can improve the robustness of species limits and validation analyses. A diverse framework linking genetic and phenotypic evidence ultimately enhances taxonomic stability and can support conservation and ecological research particularly in poorly documented or uncommon species (Padial et al. 2010; Fujita et al. 2012). Our study combined three independent lines of evidence to validate the species status of *L. minor* and objectively assess the diagnosable traits separating it from other Red Bats. Notwithstanding, we identified further questions that must be addressed to fully evaluate populations of *L. minor* across the Caribbean. These include, for example, 1. Thorough assessments of the species limits in the Bahamas where multiple Red Bats may occur in sympatry. 2. Evaluation of the range extent, density, and environmental requirements. 3. Phylogeographic analyses to document interisland gene flow or population structure. On a phylogenetic scale, obtaining proper data from *L. pfeifferi* and *L. degelidus* to examine them in a broader phylogenetic context with other Red Bats in an approach similar to what we present herein could help uncover patterns of morphological and genetic variation. These are particularly important in the broader sense of Caribbean biogeography to better understand local biodiversity, develop proper species assessments, and untangle the factors that help shape local insular communities.

## Acknowledgments

We are grateful for the legacy of decades of work on Caribbean paleofauna and contributions by C.A. Woods and other members of his field crew who collected the *L. minor* specimens used in our study. R. Hulbert at the Vertebrate Paleontology Department at UF provided access to *L. minor* specimens and processed the necessary loans. N.B. Simmons and N.P. Duncan provided access and loans to the comparative material of other Red Bats housed in the Mammalogy Collections at American Museum of Natural History. I.R. Hays kindly aided with measuring, photographing, and creating shapes from specimens. We are indebted to B. da Silva Fonseca and R.D. Barrilito for logistical support critical for the completion of this study. Work by JAS-C was partly supported by a Rutgers University Research Council Award. Work by CCA was funded by a postdoctoral scholarship at the Soto Lab of Bat Biology (SLaBB) at Rutgers University. Identification of fossil *L. minor* specimens was done by JAS-C and financially supported by a National Science Foundation (DEB-2135257) award to JAS-C.

## Supplementary Data

### Supplementary Data SD1

Database of loaned specimens examined to assess phenotypic species limits and shape descriptors of Red Bats. See file SupplementaryDataSD1.xls.

### Supplementary Data SD2

Contour traces used in our SHAPE Analysis. Four profiles were selected representing Miller’s (1931) *L. minor* diagnostic characters: A) Dorsal view to capture skull roundness; B) Posterior view, examining the level of flatness across the dorsal surface of the brain case; C) Dorsal view to assess level of development and projection of the lacrimal ridge, which was framed inside a cropping box and scaled to grid paper in the background of the photo, and oriented consistently across the profile maxillary with placement markers built into the cropping frame; and D) Lateral view of the mandible, capturing the contour of the last upper premolar PM^4^. The selected areas were converted into black and white silhouettes for analysis in the program SHAPE v1.3 (Iwata & Ukai, 2002).

### Supplementary Data SD3

Uncorrected p genetic distances estimated for 657 bp of the mitochondrial cytochrome oxidase I (COI) gene in Red Bats (*Lasiurus borealis, L. minor, L. pfeifferi*, and *L. seminolus*). Values along the diagonal represent within group genetic distances, and values below the diagonal represent between group genetic distances. Intraspecific variation not included for *L. pfeifferi* because N = 1.

